# Acute repair of meniscus root tear partially restores joint displacements as measured with MRI and loading in a porcine knee

**DOI:** 10.1101/2023.02.01.526670

**Authors:** Kyle D. Meadows, John M. Peloquin, Milad I. Markhali, Miltiadis H. Zgonis, Thomas P. Schaer, Robert L. Mauck, Dawn M. Elliott

## Abstract

The meniscus serves important load-bearing functions and protects the underlying articular cartilage. Unfortunately, meniscus tears are common and impair the ability of the meniscus to distribute loads, greatly increasing the risk for developing osteoarthritis. Therefore, surgical repair of the meniscus is a frequently performed procedure; however, this repair does not always prevent osteoarthritis. This is hypothesized to be due to altered joint loading post injury and repair, where the functional deficit of the meniscus prevents it from performing its role of distributing forces. However, many studies of meniscus function required opening the joint, which alters kinematics. The objective of this study was to use novel MRI methods to image the intact joint under axial load and measure the acute meniscus and femur displacements in an intact joint, after a meniscus root tear, and after suture repair in the porcine knee, a frequently used in vivo model. We found that anterior root tear led to large meniscus and femur displacements under physiological axial loading, and that suture anchor repair reduced these displacements, but did not fully restore intact joint kinematics. After tear and repair, the anterior region of the meniscus moved posteriorly and medially as it was forced out of the joint space under loading, while the posterior region had small displacements as the posterior attachment acted as a hinge about which the meniscus rotated in the axial plane. This technique can be applied to evaluate the effect of knee injuries and to develop improved repair strategies to restore joint kinematics.

## Introduction

The meniscus serves important load-bearing functions at the knee and protects the underlying articular cartilage. Unfortunately, meniscus tears are a common condition that impairs the ability of the meniscus to distribute loads and, without surgical treatment, greatly increases the risk for developing osteoarthritis (OA) [1]–[10]. Surgical meniscus repair is chondro-protective and can delay OA but does not halt degeneration for many patients [11], [12]. It is often hypothesized that this is due to altered joint loading post injury and repair, where the functional deficit of the meniscus prevents it from performing its role of distributing forces in the knee. Previous work has shown that there is a relation between greater meniscal extrusion and joint degeneration [13]–[15]. Some studies have shown, using pressure sensors (e.g., Tek-Scan) inserted into the joint space, that meniscus injury leads to larger peak forces and smaller contact areas on the tibial plateau; however, these studies required transecting the joint capsule and sometimes the LCL or MCL to make the measurement [2]–[4], [16], [17]. Therefore, developing non-invasive measurement of joint and meniscus kinematics is critical to understanding the impact of meniscus injury and repair on joint loading and chondroprotection.

MRI acquired under different loading conditions allows for non-invasive measurement of joint kinematics without disrupting any joint tissues. While forces cannot be measured via MRI, kinematics, including displacements and rotations of the meniscus and femur, can be measured in the intact joint. This is important as the collateral ligaments and joint capsule contribute meaningfully to joint kinematics. Two studies have utilized MRI to measure meniscus protrusion/extrusion at various flexion angles in healthy, uninjured subjects or in patients with OA, but not following a diagnosed meniscus injury [18], [19]. Other work used MRI to measure 3D strains in the meniscus at various degeneration levels, but either did not report femur or meniscus displacements or only studied healthy joints [20]–[22]. Additional work has measured cartilage deformations under loading using MRI-compatible loading devices or exercise, but meniscus findings were not reported [23]–[25]. Evaluation of meniscus repair methods would benefit from measurement of joint kinematics after surgery in an intact joint. Likewise, application of novel imaging protocols, for example T1 VIBE MRI, that allow for better imaging of the menisci than traditional clinical MRI used in other studies, would improve our understanding of meniscus function under load. Thus, the objective of this work was to measure changes in meniscus and femur displacements in an intact joint, following a meniscus root tear, and following suture repair at an acute time point using novel MRI methods to image the joint under axial load. We hypothesized that a root tear would cause meniscus extrusion and medial displacements of the femur, and that a suture anchor repair would restore the meniscus to its near intact position and kinematics. This in vitro work used a Yucatan minipig stifle joint model, similar to our established animal model [3], [4].

## Methods

To measure the effect of injury and repair on meniscus and femur displacements with loading, each joint was loaded and imaged in three experimental states: Intact, Tear, and Repair.

### Sample Preparation

Seven fresh-frozen Yucatan minipig hindlimbs, harvested from 12–18 month-old castrated males, were obtained (Sierra Medical for Sciences, Whittier, CA, sierra-medical.com). We chose this age range because the animals are skeletally mature and it matches the age of pigs that we have previously used in our live animal model [3], [4]. After thawing, all skin and musculature were removed from the stifle joint, with special care taken to preserve the joint capsule, lateral collateral ligament, and medial collateral ligament. The femur and tibia were then cut approximately 4 inches from the joint line to obtain an intact joint. From this point forward, the joint was maintained in PBS-soaked gauze to avoid dehydration.

The femur and tibia were potted in Ortho-Jet acrylic resin (Polymethyl Methacrylate, PMMA, from Lang Dental, Wheeling, Illinois) using Delrin molds for gripping in an MRI-compatible custom-built loading frame (Fig 1A). The embedded tibia is secured via set screws in a cup attached to the bottom of the frame, and the embedded femur is secured in a cup attached to a threaded rod that slides freely through a hole in the top of the frame. The displacement of the rod and hence the femur can be fixed in place via lock nuts. For this study, the tibia cup was angled to produce 30° joint flexion to match previous studies and because this is approximately maximum extension in the pig, representing 0° flexion in a standing human [21], [26]. Fluoroscopy was used to adjust and confirm joint alignment, including parallel alignment of the tibia and femur with the molds.

**Figure 1:**
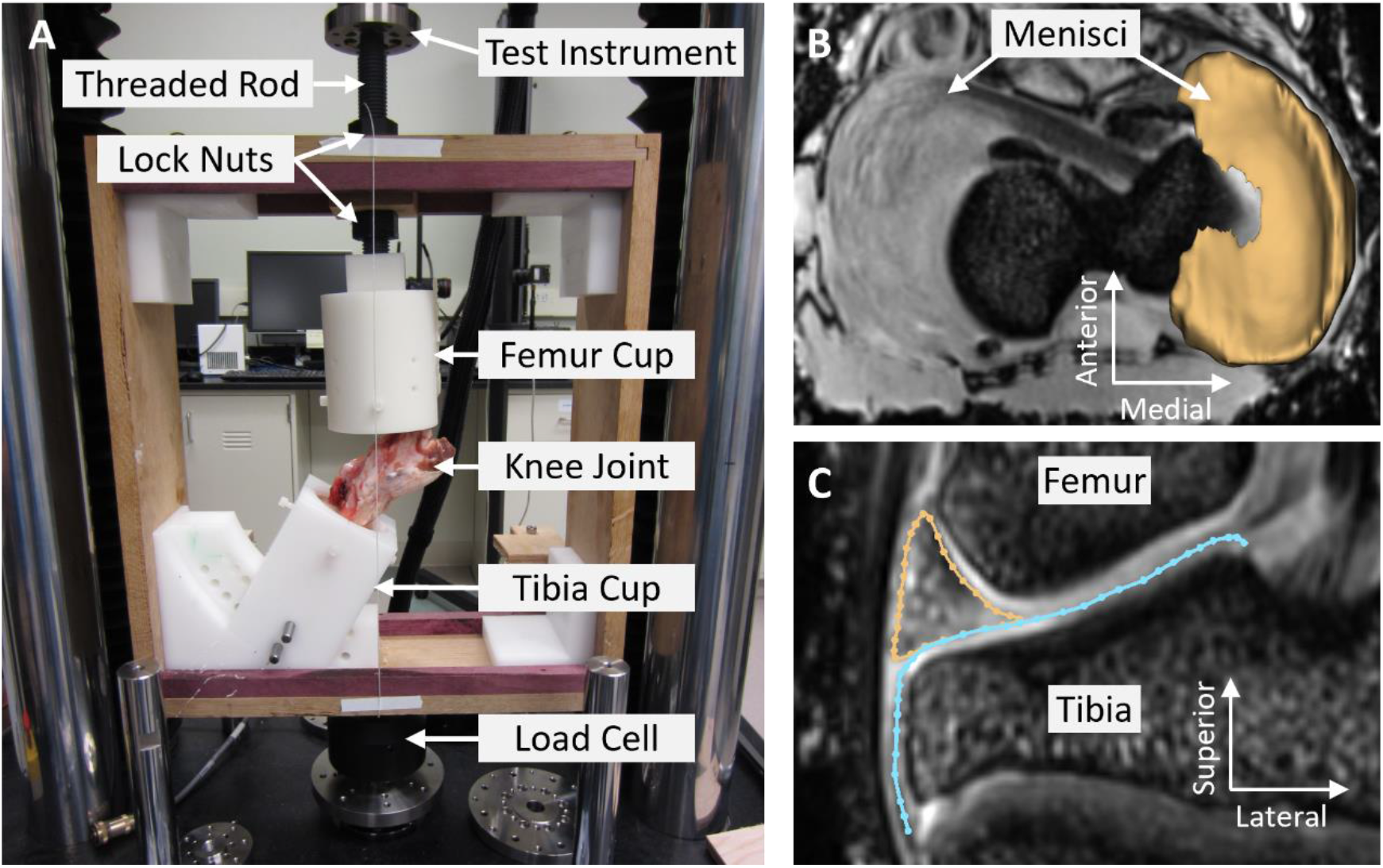
A) MRI compatible loading frame in the mechanical testing instrument (TA ElectroForce). T_1_ VIBE MRI of knee in (B) axial plane with 3D segmentation of medial mensicus overlayed, and (C) mid-coronal plane with outlines of meniscus (yellow) and tibial plateau (blue) overalyed.

The Intact case was an intact joint, where the joint capsule and lateral and medial collateral ligaments were preserved and the musculature removed, prepared as described above. In the Tear case, a medial meniscus root tear was created by transection of the anterior attachment (destabilization of the medial meniscus, DMM, a frequently used model) using a minimally invasive anterior approach to limit the size of the incision in the joint capsule [3], [4], [10]. The attachment was severed via a horizontal cut along the bone insertion. Repair was done by an experienced veterinary surgeon (TS) using a SwivelLock suture anchor (Arthrex) placed into the bone such that the meniscus was restored to its original Intact position via sutures running from the anchor through the anterior attachment. This is a technique used for root tear surgical repair in humans [27]–[29].

### Joint Loading and MRI

To non-invasively measure joint kinematics, MR images were acquired at two loading states. For this experiment, we used a preload (Low Load) of 44.5 N (10 lbf) and a High Load of 850 N or approximately 1.5× body weight (15-month-old male Yucatan weighs ∼60 kg). The Low Load ensured the joint was in contact and was applied by placing a mass on the upper nut on the threaded rod, then locking in the displacement using the bottom nut. For High Load, the joint was loaded using a mechanical test instrument (TA ElectroForce 3510, TA Instruments, New Castle, DE) pressing on the threaded rod. Load was applied by manually moving crosshead at approximately 0.05 mm/s until the target 850 N load was reached, holding displacement for 10 s to allow the joint to relax, and reapplying the target load. The nuts on the threaded rod were then tightened until zero load was read by the test instrument’s load cell, transferring all load to the loading frame and locking in the applied displacement. Note the load can relax further even with the lock nuts tightened due to viscoelasticity, potentially producing motion within the joint articulation or within tissues. Thus, load was consistently applied 30 min prior to MRI scanning to avoid motion during scanning.

MRI was performed on each joint after Low Load and High Load using the same protocol. Two 4-channel surface coils were wrapped around the joint using Velcro, and a T^1^ VIBE sequence was run to image the joint at high resolution (TR = 10 ms, TE = 3.45 ms, voxel size = 0.2 mm x 0.2 mm x 0.2 mm, runtime = 18 min, Fig 1B-C). This sequence was adapted from bone MRI, and new parameters used here were developed in pilot studies by optimizing the sequence for meniscus signal intensity [30], [31]. After High Load imaging, residual (unrelaxed) load was measured by remounting the loading frame in the TA ElectroForce 3510 and releasing the lock nuts. Residual loads were 295N ± 41N at 68 min ± 4 min after the load was initially locked into the frame (∼65% relaxation from initial load to the end of imaging). These values were not different between treatment groups (p=0.68, ANOVA, n=4).

Each joint was loaded and imaged in three different experimental states: Intact, Tear, and Repair. After initial loading and MRI in the Intact state, joints were refrigerated at 4°C overnight to equilibrate under zero applied load. The next day, the Tear case was created via release of the anterior attachment 30 minutes before loading, and loading and MRI were repeated using the same protocol as above. Joints were frozen after Tear MRI so that Repairs could be done as a batch. Loading and MRI protocols were repeated on the Repaired joints either on the same day as the repair procedure or the following day after overnight refrigeration.

### MRI Image Analysis

Displacements of the medial meniscus and femoral condyles in the Intact, Tear, and Repair conditions were measured from the MRI images relative to the Intact Low Load position. First the Intact Low Load image was resliced into the scanner RAS+ (Right, Anterior, and Superior positive) coordinate system, with 0.2 mm isotropic voxels. Next, a tibia-aligned coordinate system was defined. Then, a partial segmentation of the tibia was made using ITK-SNAP’s active contour semi-automatic segmentation tool [32]. The tibia coordinate system and segmentation were created in the Intact Low Load image, and the five other images (Intact High Load, Tear Low and High Load, and Repair Low and High Load) were aligned to this coordinate system by rigid registration of the tibia, using the segmentation as a registration mask [20], [21], [33]. All registrations were visually checked to confirm alignment of the tibia across all 6 conditions. All meniscus and femur displacements were calculated in tibia-aligned coordinates with medial, anterior, and inferior assigned as the positive directions.

To measure meniscus displacement, the outer edge of the medial meniscus was traced in 3D following a line on the outer surface halfway between the superior and inferior rims. This labeling was done for all conditions, in the tibia aligned coordinate system. Displacements were measured from Intact Low Load to each High Load condition by matching points at the same % arc length along the traced outer edge lines and subtracting their positions. To measure the change in starting position due to treatment, displacements were also measured in the same way for Tear and Repair Low Load relative to Intact Low Load. For regional analysis, the points comprising the outer edge line and their associated displacement values were grouped into equal thirds: the anterior region (AR), mid-body region (MR), and the posterior region (PR).

Displacements of the medial and lateral femoral condyles were measured at the points where the condyles contacted the tibial plateau in the Intact Low Load image. These contact points were labeled using 3D Slicer in the aforementioned tibia-aligned coordinate system [34]. The femur was then automatically segmented, and the segmentation used as a mask for rigid registration of the femur between the Intact Low Load condition and the other five conditions. The resulting rigid transform matrices were applied to the Intact Low Load positions of the two condylar contact points to calculate their displaced positions. Displacements were then calculated by subtracting the positions from Intact Low Load to each other condition.

Statistical analyses used a repeated measures ANOVA to test for the effect of treatment (Intact, Tear, Repair) on the meniscus and femoral condyle displacements using JMP 16 (JMP, Cary, NC). If significance was met, a post-hoc Tukey test was carried out to test for differences between treatment groups. Pearson’s correlations were performed to test for relationships of the displacements between regions and between the meniscus and femoral condyle. Significance was set at p<0.05.

## Results

### Meniscus Displacements

Qualitatively, the midcoronal outline (Fig 2A) and outer boundary (Fig 2B) of the meniscus under loading revealed that the meniscus was extruded outside of the joint space following Tear. Repair partially restored the original position of the meniscus within the joint space. In all treatment groups, High Load caused the meniscus to partially extrude from the joint space, consistent with uninjured human studies (Fig 2A,B) [18], [19]. The mid-coronal slice was defined in the Intact Low Load case, then the same slice was traced for all other conditions. The largest Intact Low Load → Treatment High Load displacements occurred in the anterior meniscus, which moved medially out of the joint space, whereas the posterior meniscus showed little displacement, probably due to the still-intact posterior attachment (Fig 2B). Based on this, the meniscus displacements were averaged within three regions of interest: anterior region (AR), mid-body region (MR), and posterior region (PR) (Fig 2C). Quantitative analyses supported these observations, as shown by the displacement vectors from Intact Low Load to Intact High Load (Fig 2C), Tear High Load (Fig 2D), and Repair High Load (Fig 2E) for a representative sample. For the AR, at Intact High Load, displacements were small and mostly uniform (Fig 2C). However, following Tear and Repair there were large medial displacements at this location (Fig 2D,E). In addition, small anterior displacements in the Intact case changed direction for Tear and Repair and were posterior in both the AR and MR regions. In contrast, the PR had small displacements in all conditions (Fig 2C-E).

**Figure 2:**
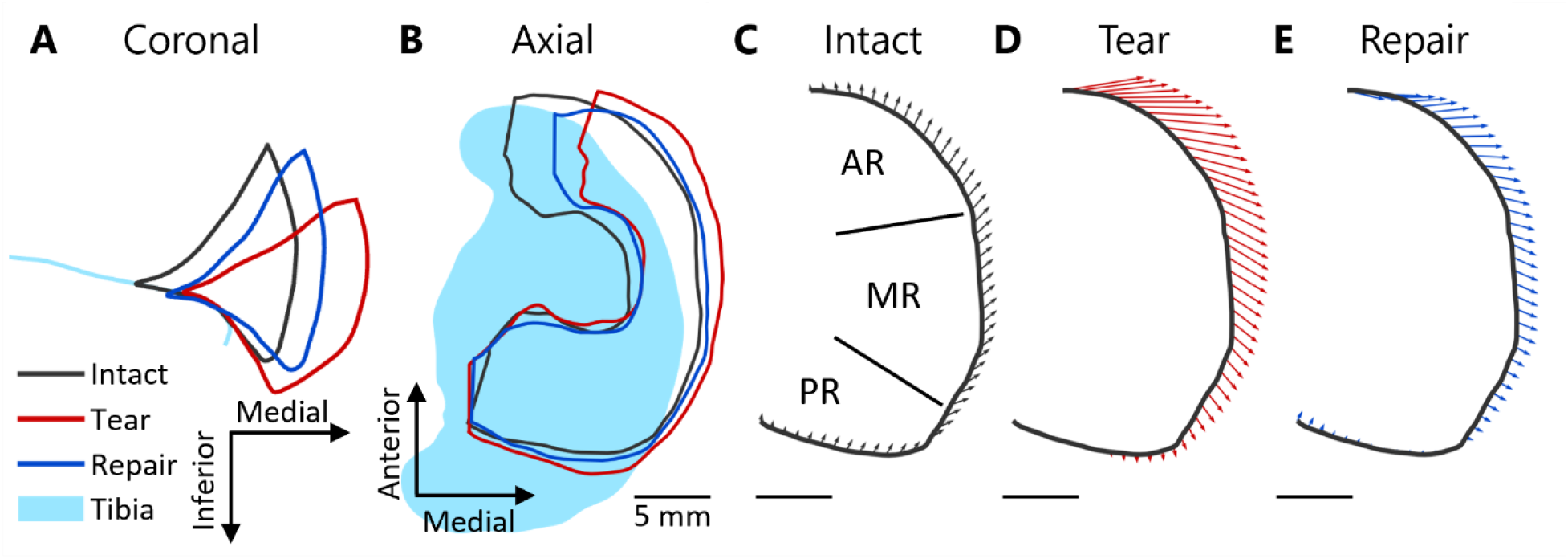
Representative example of meniscus displacements at High Load with each treatment. (A) Mid-coronal outlines show that the Tear allowed the meniscus to be pushed out of the joint space, while the Repair partially restored the meniscus position towards the Intact case. (B) Axial plane outlines show the same, while also illustrating that the anterior part of the meniscus displaced more than the posterior. (C) Region labels (AR = Anterior Region, MR = Mid-Body Region, PR = Posterior Region) for a representative sample of displacements from Intact Low Load to Treatment High Load for (C) Intact, (D) Tear and (E) Repair treatment. The anterior region of the meniscus had large medial displacements following Tear that were reduced with Repair. The MR had moderate displacements that increase with Tear and moved posteriorly, while the PR had small displacements.

Displacement vectors of the outer meniscus boundary in each region were quantified under High Load relative to the Intact Low Load (Fig 3A-C). For statistical analyses, we isolated each vector component (anterior and medial directions) (Fig 3D-I). The AR and MR had significant effects of treatment in both directions (p<0.05), but the PR only had a significant effect of treatment in the anterior direction (p=0.02 for anterior direction and p=0.06 for medial direction), post-hoc statistics are described below.

**Figure 3:**
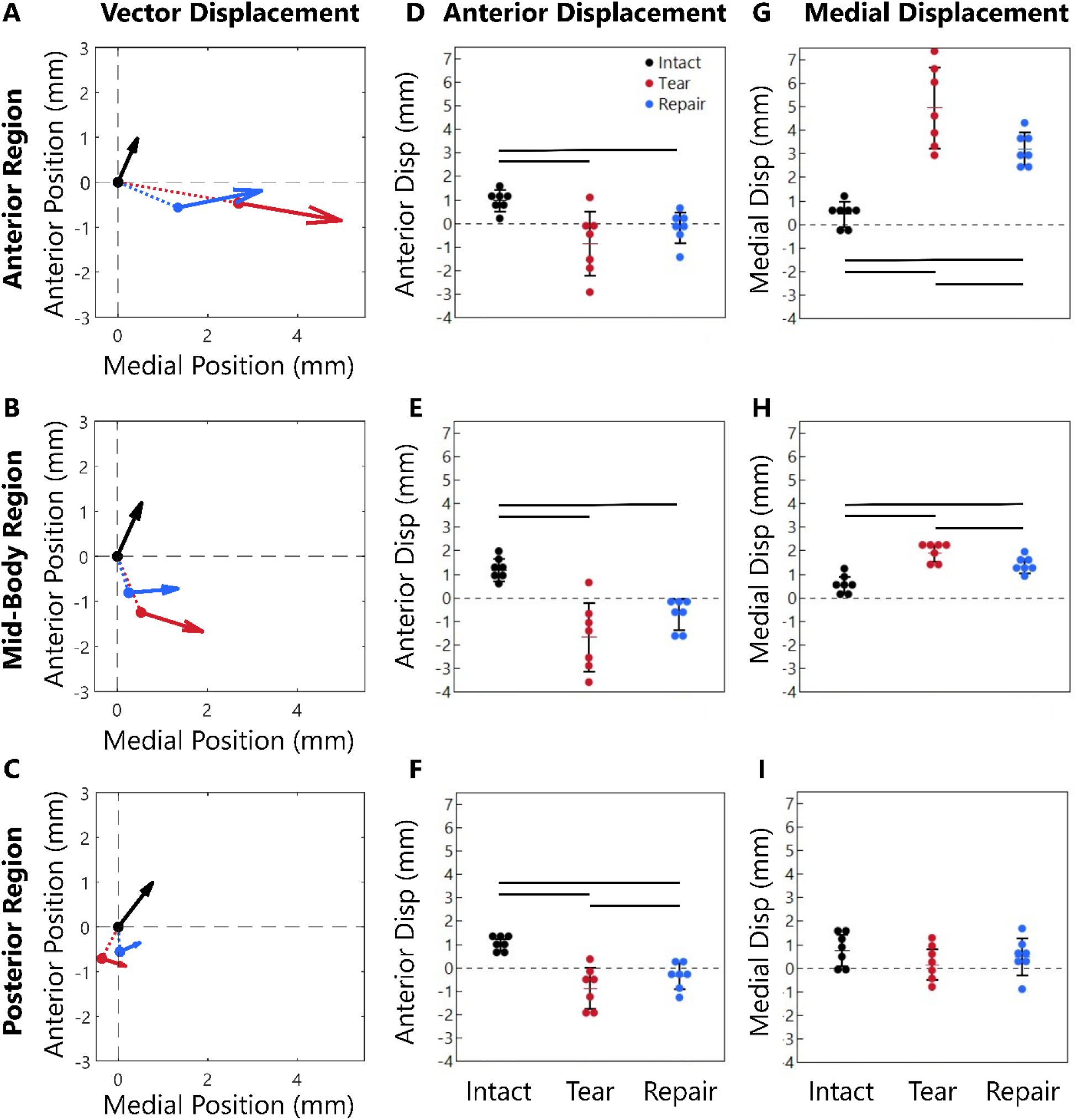
Meniscus position at High Load (arrow head) and Low Load (arrow origin) in the (A) Anterior Region (AR), (B) Mid-Body Region (MR), and (C) Posterior Region (PR). The dashed lines in A-C show displacement from the Intact Low Load position to the Tear and Repair Low Load initial positions (additional information provided in supplemental data). The total displacement vector for each case, Intact Low Load to Intact/Tear/Repair High Load, is decomposed into (D-F) anterior displacement components and (G-I) medial displacement components. All meniscus regions displaced posteriorly after the Tear (D-F) and the AR and MR displaced medially (G-I). The Repair generally reduced the large displacements induced by the Tear but did not completely restore the Intact displacements.

For the Intact case, the AR moved 0.97 mm anteriorly and 0.43 mm medially (Fig 3D,G). With a Tear, the AR moved 0.85 mm posteriorly (similar magnitude but in the opposite direction compared to Intact, p<0.05) and 4.96 mm medially out of the joint space (11.5× Intact, p<0.05, Fig 3D,G). With Repair, the AR displacements were smaller, moving only 0.19 mm posteriorly (p<0.05) and 3.19 mm medially (7.4× Intact, p<0.05). While the Repair reduced the meniscus displacement, it did not return to Intact levels. The vector visualization shows that in the AR the Tear caused large medial displacements and small posterior displacements, and that the medial displacements were partially restored with Repair (Fig 3A).

The meniscus displacement changes were similar in the MR compared to the AR, but with lower magnitudes overall (Fig 3E,H). For the Intact case, the MR displaced 1.19 mm anteriorly and 0.54 mm medially, and with the Tear the MR displaced 1.66 mm posteriorly (opposite direction from Intact, p<0.05) along with 1.89 mm of medial displacement (3.5× Intact, p<0.05). With Repair the displacement again decreased compared to Tear, but the MR still displaced 0.72 mm posteriorly (opposite of Intact, p<0.05) and 1.34 mm medially (2.5× Intact, p<0.05). The MR moved more posteriorly and less medially than the AR. The vector visualization demonstrates that in the MR, the Tear caused large posterior displacements that were partially restored with Repair, and that the medial displacements were overall smaller than in the AR region (Fig 3B).

The PR had the smallest displacement in both directions, with statistical significance between treatments only in the anterior direction (p=0.02 for anterior direction and p=0.06 for medial direction). The Intact PR only displaced 0.99 mm anteriorly and 0.76 mm medially (Fig 3F,I). With a Tear, the PR displaced 0.88 mm posteriorly (opposite direction from Intact, p<0.05), but little medially (0.16 mm, 0.2× Intact). With the Repair the displacements were more similar to Intact, with 0.36 mm posteriorly (opposite direction from Intact, p<0.05) and 0.48 mm medially (0.6× Intact). The vector visualization demonstrates that, in the Intact case, the PR displacements are similar to the AR and MR, but with both Tear and Repair, PR displacements were much smaller (Fig 3C).

We next evaluated correlations between anterior and medial displacements and between regions. Anterior and medial displacement components were strongly correlated in the AR (R^2^= 0.85, p<0.05, Fig 4A) and the MR (R^2^=0.52, p<0.05) but not in the PR (R^2^= 0.11, p=0.15). With high medial displacements, the meniscus moved posteriorly. In contrast, low medial displacements, as in the Intact case, caused anterior meniscus displacements. In terms of comparison between regions, the anterior displacement component was strongly correlated between all regions: AR and MR (R^2^=0.88, p<0.05, Fig 4B), AR and PR (R^2^=0.75, p<0.05), MR and PR (R^2^=0.90, p<0.05). For the medial displacement component, there was a significant relationship only between AR and MR (R^2^=0.70, p<0.05, Fig 4C), while the AR to PR (R^2^=0.08, p=0.22) and MR to PR (R^2^=0.00, p=0.94) had no relation, likely due to the small displacement magnitudes in the PR.

**Figure 4:**
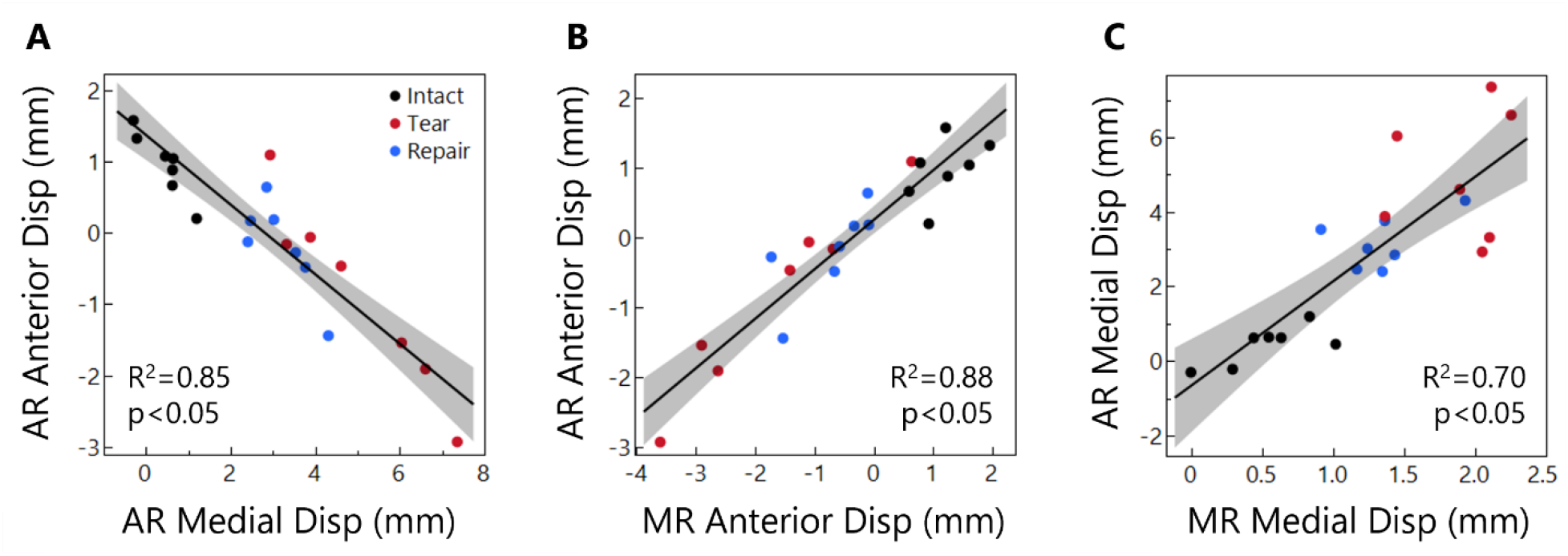
(A) Relationship between AR anterior and medial displacement. Greater medial displacement caused posterior displacement. (B) Relationship between AR and MR anterior displacement. (C) Relationship between AR and MR medial displacement.

### Femoral Condyle Displacements

In addition to meniscus displacements, we also evaluated the 3D displacements of the medial and lateral femoral condyles at their contact point with the tibial plateau (Fig 5). For the medial condyle (Fig 5A), Intact Low Load → Treatment High Load displacements were in the anterior, medial, and inferior directions for all treatments. Differences between treatment groups were non-significant, but barely so (p=0.06 anterior direction, p=0.08 medial direction, and p=0.11 inferior direction). Anterior displacement of the medial condyle was 2.76 mm for the Intact case. This was reduced to 2.03 mm with Tear but returned to 2.53 mm with Repair, nearly the Intact value. Medial displacement in the Intact case was 0.42 mm, which was 3.5× larger with Tear (1.54 mm). With Repair, medial displacement was partially restored (1.23 mm), but was still 3× larger than Intact. In the inferior direction, the medial condyle displacement was 1.26 mm for Intact. In the Tear case, the inferior displacement was slightly larger (1.40 mm) and was even larger following Repair (1.51mm). The lateral condyle, similar to the medial condyle, displaced in the anterior, medial, and inferior directions for all treatments, and there were no significant differences with treatment (p=0.08 anterior direction, p=0.08 medial direction, and p=0.09 inferior direction). Lateral condyle displacement results are provided in the supplemental section. Overall, despite lack of significance, these results suggest that severing the anterior medial meniscus attachment (Tear group) altered the displacement of the femur relative to the tibia, and Repair partially restored the kinematics towards that of the Intact condition.

**Figure 5:**
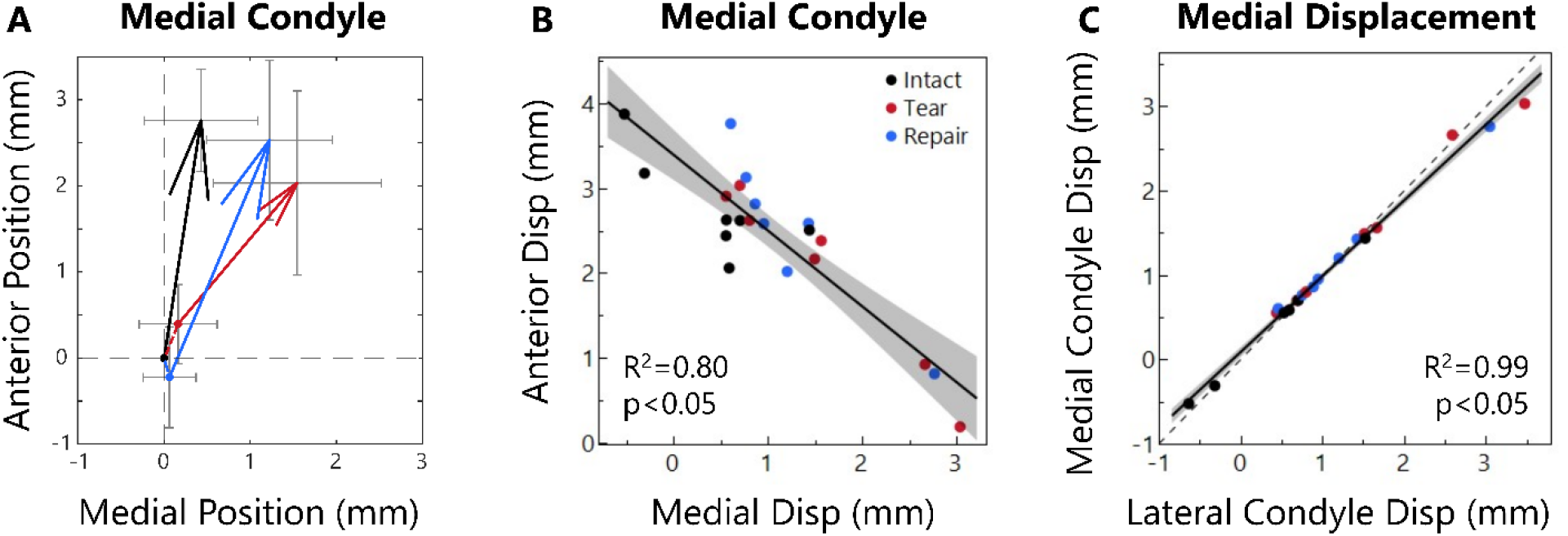
(A) Medial condyle positions at Low Load (arrow origin) and at High Load (arrowhead). Intact joints move anteriorly about 2.76 mm and medially < 0.5mm. The Tear case allowed the contact points to shift only 2.03 mm anteriorly but 1.54 mm medially. Repair made motion more similar to Intact with 2.53 mm anterior and 1.23 mm medial displacement. (B) Relationship between Medial Condyle Anterior and Medial Positions. As the condyle moves further medially, it moves less anteriorly. (C) Medial vs Lateral condyle displacement in the medial direction. The relation is nearly perfectly 1:1 (dashed line).

There was a strong correlation between the medial condyle’s anterior and medial displacement components (R^2^=0.80, p<0.05, Fig 5B), similar to the meniscus. While there was a low correlation between the anterior and inferior displacement (R^2^=0.27, p<0.05), there was a moderate correlation between the medial and inferior displacement (R^2^=0.47, p<0.05), likely related to the slope of the tibial plateau. The lateral condyle exhibited similar correlations between displacement components. As expected for a rigid body, displacements of the medial and lateral condyle were significantly correlated in all directions, including anterior (R^2^=0.41, p<0.05), medial (R^2^=0.99, p<0.05, Fig 5C), and inferior (R^2^=0.60, p<0.05).

### Correlations between Meniscus and Femur Displacement

Finally, we evaluated correlations between the meniscus regions and the medial femoral condyle displacements. Considering the meniscus AR, the anterior displacement components of the AR and femoral condyle were correlated (R^2^ = 0.57, p<0.05, Fig 6A). In the Tear and Repair groups, the meniscus moved posteriorly while the medial condyle moved anteriorly. In the Intact case, however, the meniscus moved anteriorly, and the medial condyle moved anteriorly to a greater degree. The medial displacement components of the AR and the medial condyle were also highly correlated (R^2^ = 0.65, p<0.05, Fig 6B). In the Tear and Repair groups, the meniscus moved much further medially than the condyle (up to 7.4 mm vs up to 3.0 mm, respectively), however, the Intact group lies mostly along the 1:1 line where meniscus and condyle motion were equal.

**Figure 6:**
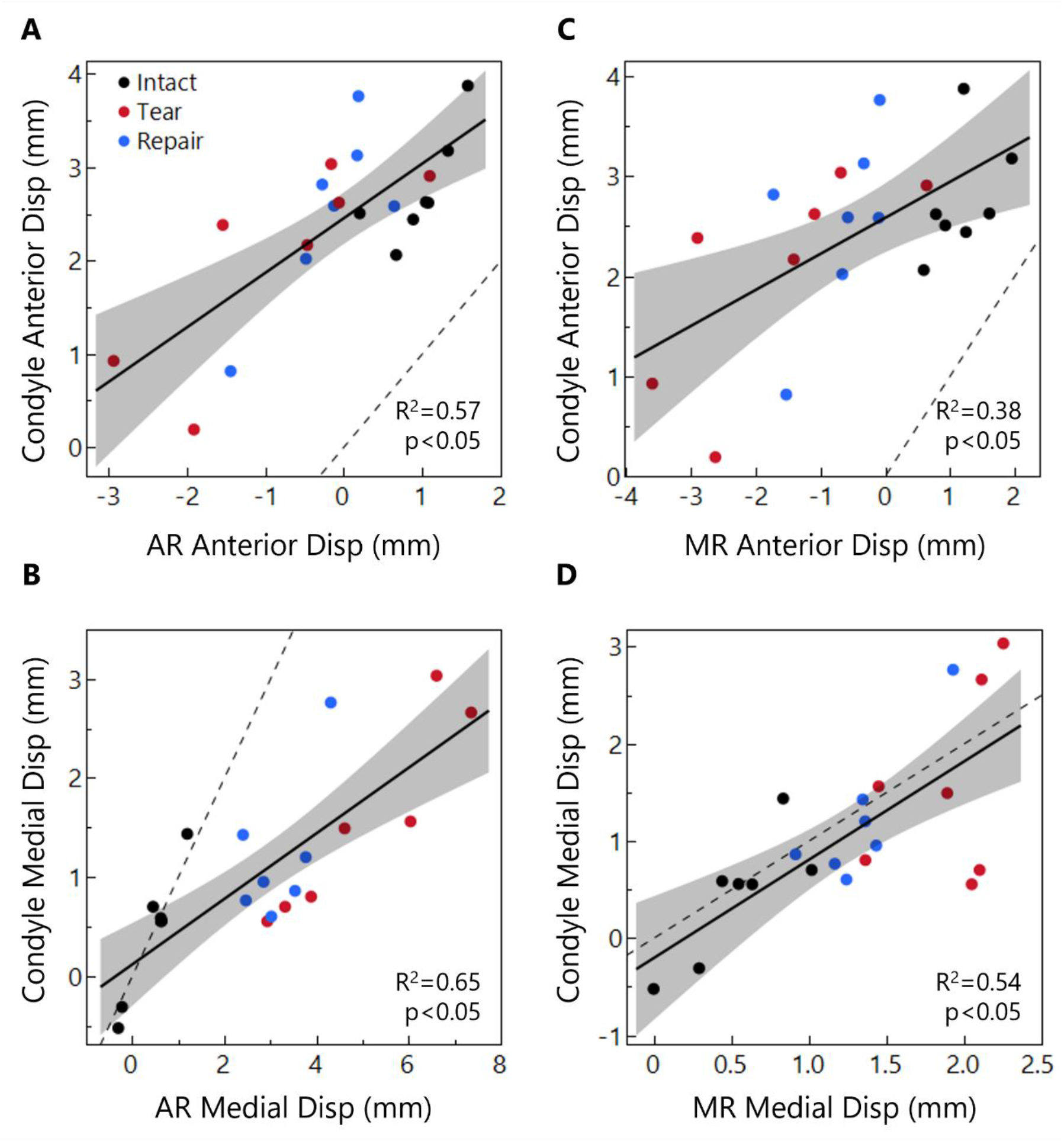
Displacement relationships between the medial condyle contact point and the meniscus AR (A, B) and MR (C, D), considering anterior (A, C) and medial (B, D) displacement components. The condyle moves further anteriorly than the meniscus (A, B), while medial displacements (B, D) are closer to a 1:1 relationship (dashed line) between the tissues. This 1:1 relationship is lost in the AR with Tear and Repair conditions because with the anterior medial meniscus attachment released, the meniscus AR displaces far outside of the joint space with loading.

Considering the meniscus MR, the anterior displacement components of the MR and medial condyle were significantly correlated (R^2^=0.38, p<0.05, Fig 6C). Menisci in the Intact group moved anteriorly, but in the Tear and Repair groups, the MR moved posteriorly. The medial displacement components of the MR and the femoral condyle were not correlated as strongly as in the AR, but in contrast to the AR region their relationship followed the 1:1 line very closely, with only slightly more meniscus displacement than contact point displacement (R^2^ = 0.54, p<0.05, Fig 6D). There were no significant relationships between PR meniscus displacement and femoral condyle displacement.

## Discussion

In this study, we measured joint kinematics in an intact porcine knee, after anterior root tear, and after repair using non-invasive MRI to maintain an intact joint capsule and surrounding ligaments. We found that transection of the medial meniscus attachment (Tear) allows for large amounts of medial extrusion of the medial meniscus, particularly in the anterior region. A suture anchor repair partially restores the meniscus in the joint space and limits femur displacement but does not fully restore kinematics to match the Intact case. This study was performed using an MRI-compatible loading device and a newly adapted MRI sequence allowing us to capture the joint structure at multiple loading conditions and with much higher meniscus signal and resolution than previously reported.

When comparing the regions of the meniscus, the AR had the most displacement, while the PR had the least displacement. The combined effect in the Tear and Repair cases was to rotate the meniscus in the axial plane; the AR and MR move medially, while the PR remains mostly static due to its posterior attachment (Fig 2B, 3). The posterior attachment behaves like a hinge about which the meniscus rotates. This effect also explains the differences between displacement of the meniscus and of the femur. Though the femur moves anteriorly in all cases due to the 30° flexion angle, it moves less anteriorly and more medially after Tear and Repair, pushing the meniscus AR out of the joint space (Fig 6). With extrusion, the meniscus pivots about the posterior attachment, and the AR and MR regions of the meniscus move slightly posteriorly. Similar work found that the meniscus of an intact joint instead moved posteriorly with load [21]. The likely causes of this difference in findings is that our study applied load along the femoral axis, pushing the femur forward over the tibia, while the other study applied load along the axial direction of the tibia, pushing the femur backward over the tibia.

Our findings for the femur displacements agree with previous studies of joint kinematics. Studies using pressure sensors placed into the joint space showed that the femoral contact point shifts medially after meniscus injury [2]–[4], [16], similar to our meniscus and femur results. Contact pressure tracking primarily finds that the contact area decreases, indicating that the meniscus is not carrying load due to extrusion by the femur. This matches what we find when we track both the femur and meniscus in our study. After Tear, the femur moves medially and anteriorly, pushing the meniscus, especially the anterior region, out of the joint, so that the meniscus is no longer carrying load or distributing forces to protect the cartilage. This is likely linked to the higher rates of OA seen in many patients with high meniscus extrusion or after meniscus root tear [13]–[15].

The large displacements in the Repair case, particularly in the AR, suggest that a better surgical treatment might be required to restore joint kinematics to the Intact state. The suture anchor repair returns the anterior aspect of the meniscus back to its natural position by suturing through the cut anterior attachment. Due to the highly aligned fiber structure of the attachment, it is possible that the anchor did not adequately hold the meniscus in place due to the sutures pulling though the weaker inter-fiber matrix when load was applied. However, the present study explored the effectiveness of Repair treatment at acute time points, whereas in vivo, there would be time for healing, remodeling, and potentially unloaded post-operative care. Thus, further in vivo studies are needed to explore if the suture anchor repair aids in healing to increase the strength of the anchor/anterior attachment complex and to reduce medial extrusion of the meniscus under load. Another solution to reduce medial extrusion may be to add sutures on the outside of the joint for meniscus centralization, especially in the anterior half, similar to studies by the Sekiya group [15], [16]. The methods used in the present study are valuable to test other repair strategies to restore joint kinematics and meniscus load distribution to protect the underlying cartilage.

Our novel T1 VIBE MRI sequence for joint imaging allowed for excellent meniscus signal intensity and resolution. Due to the meniscus’ densely packed fiber network, MRI signal in most clinical studies and cartilage-focused studies is quite low, such that in most clinical and pre-clinical research using MRI, the meniscus is a black object between the femur and tibia. With our MRI sequence, we could observe meniscus structure for segmentation and calculation of displacements. T_1_ VIBE has traditionally been utilized in abdominal, breast, and brain imaging, but has recently been recommended for use in musculoskeletal applications as a replacement for CT due to its ability to provide high resolution bone imaging [30], [31]. Prior to this study, in preliminary work, we optimized the T_1_ VIBE sequence parameters for meniscus imaging. To our knowledge, we are the first to utilize T_1_ VIBE in the knee specifically for meniscus imaging. There is a clear improvement in our images and images from similar studies to achieve more signal from the meniscus while maintaining higher resolution and/or shorter scan times [13], [14], [20]–[22].

This study is not without limitations. First, though we kept the joint capsule and surrounding ligaments as intact as possible, a small incision was required for creation of the tear and for the repair procedure. This was an approximately 3.0 cm vertical incision in the joint capsule to the medial side of the patellar tendon, which is considerably smaller than the incisions or dissections used in other studies using pressure sensors [2]–[4], [16], [17]. Secondly, due to the aligned nature of the meniscus attachment, there is the possibility of suture slippage or pull-out in the Repair group. Despite this possibility, after testing, the repair was inspected to confirm that the meniscus was still anchored to the bone as intended. The repairs remained in place, with at least 30 N of tensile load-bearing capacity.

In conclusion, this study found that anterior root tears resulted in large meniscus and femur displacements with loading, and that suture anchor repair reduced these displacements, but did not restore intact joint kinematics. The femur moved anteriorly after Tear and Repair. Under load, the anterior region of the meniscus moved posteriorly and medially as it was forced out of the joint space, while the posterior region had small displacements due to the restraint of the posterior attachment, which acted as a hinge about which the meniscus rotated in the axial plane. The experimental techniques used here can be applied to evaluate the effect of knee injuries and to develop improved repair strategies to restore joint kinematics after injury.

## Supplemental

### Initial Positions at Low Load – Meniscus

While the displacements from Intact Low Load to Treatment High Load’s final positions are informative for overall joint kinematics, the shift in initial position between Intact Low Load and Tear/Repair Low Load is also potentially significant. Releasing the anterior attachment and repairing it alters the natural joint position of the meniscus, even at very low loads. The initial position of all three meniscus regions (AR, MR, and PR) at Low Load was measured for the Tear and Repair cases, the same way as their final positions at High Load were measured, and used to calculate the displacement from Intact Low Load to Tear/Repair Low Load (Figure, below). This shift in Low Load position primarily occurred in the AR where the Tear caused a 0.46 mm posterior and 2.69 mm medial shift in initial position. The Repair again limited this shift to 0.56 mm posteriorly and 1.33 mm in the medial direction. Both were significantly different from Intact in the medial direction, but only Repair was different in the anterior direction. The MR had significant shifts in initial position in both anterior and medial directions for both Tear and Repair where Tear shifted 1.25 mm posteriorly and 0.53 mm medially and Repair shifted 0.81 mm posteriorly and 0.25 mm medially (p<0.05). The PR had a significant shift in the anterior direction, but not in the medial direction, for both Tear and Repair. Tear shifted 0.71 mm posteriorly and 0.37 mm laterally and Repair shifted 0.56 mm posteriorly and 0.03 mm medially (p<0.05). Across the board, the Repair limited the change in initial position compared to Tear but failed to restore Intact positions.

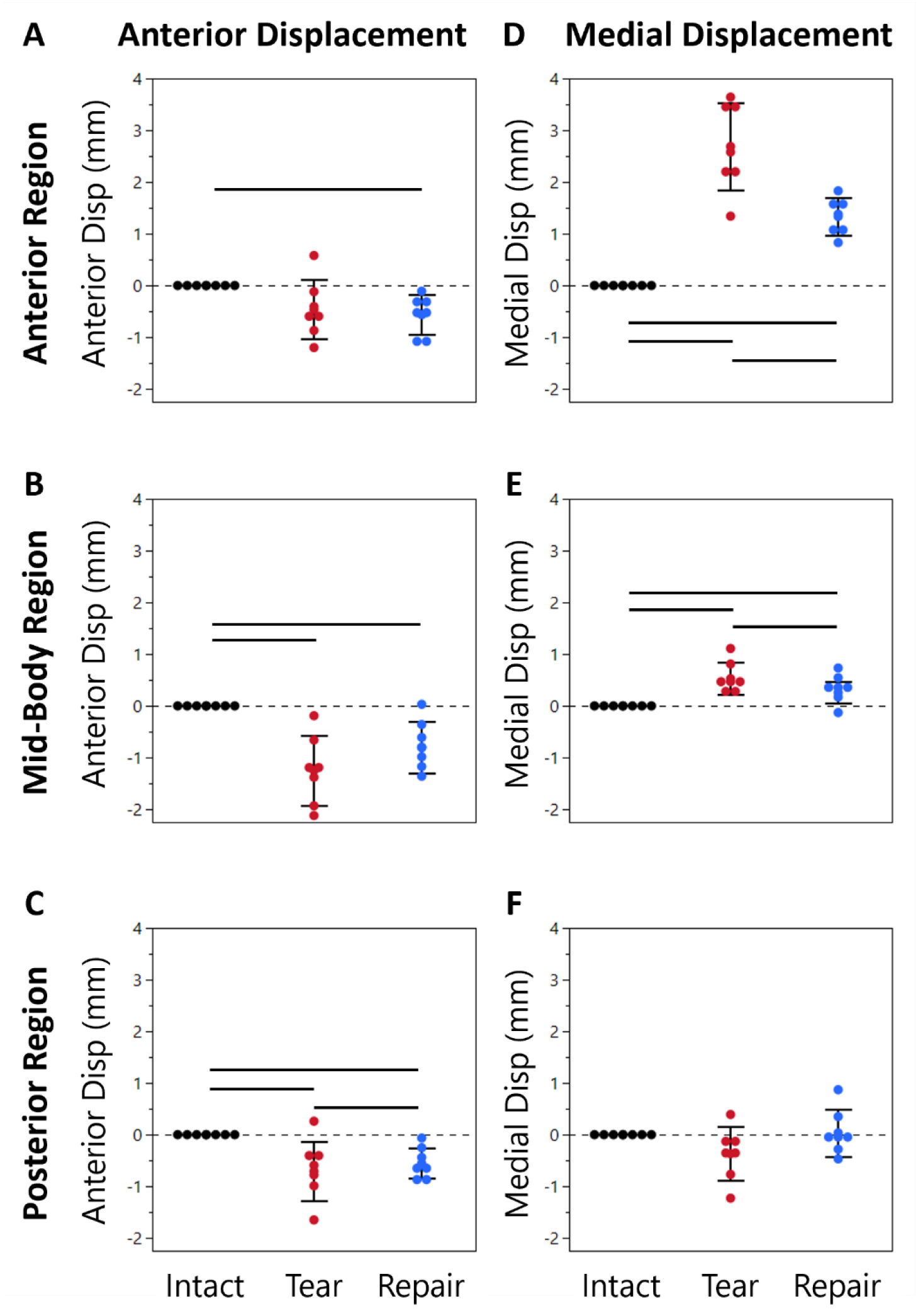

### Lateral Condyle Displacement

Lateral condyle displacements reported here are from Intact Low Load to Intact/Tear/Repair High Load. In the anterior direction, the Intact displacement was 2.60 mm. The anterior displacement was larger with Tear and Repair (2.96 mm and 3.02 mm respectively). In the medial direction, the Intact displacement was 0.42 mm. With Tear, the condyle displaced 3.8× Intact (1.60 mm). With Repair, the condyle displaced only 3.0× Intact (1.25mm). Finally, in the inferior direction, for the Intact case the condyle displaced 1.62 mm. With Tear, the lateral condyle displaced only 1.49 mm, but with Repair, the lateral condyle displaced 1.72 mm.

### Initial Positions at Low Load – Condyles

The initial positions of the femoral condyles at Low Load were significantly shifted, relative to the original Intact Low Load position, for the Tear case but not for the Repair case. The Tear case caused the initial position of both condyles to shift anteriorly (medial: 0.40mm – p=0.06, lateral: 0.69mm – p<0.05) and inferiorly (medial: 0.46mm – p<0.05, lateral: 0.42mm – p<0.05) relative to the intact case. The condyles also shifted approximately 0.2mm medially but this was not significantly different from intact. The Repair case resulted in much smaller shifts in the initial position of the condyles that were not significant. The largest shift was the medial condyle moving 0.22mm posteriorly. The medial shift was <0.13mm.

## Acknowledgements

We like to thank our funding sources for their contributions to this work.

